# Monocarboxylate transporter antagonism reveals metabolic vulnerabilities of viral-driven lymphomas

**DOI:** 10.1101/2020.12.04.410563

**Authors:** Emmanuela N. Bonglack, Joshua E. Messinger, Jana M. Cable, K. Mark Parnell, James Ch’ng, Heather R. Christofk, Micah A. Luftig

**Affiliations:** Department of Pharmacology and Cancer Biology, Duke University School of Medicine, Durham, NC, 27710; Department of Molecular Genetics and Microbiology, Duke Center for Virology, Duke University School of Medicine, Durham, NC, 27710; Active Motif, Inc., 1914 Palomar Oaks Way # 150, Carlsbad, CA 92008; Vettore, LLC, San Francisco, CA, 94158; Department of Pediatrics, Division of Hematology/Oncology, David Geffen School of Medicine at UCLA, Los Angeles, CA, 90095; Department of Biological Chemistry, David Geffen School of Medicine at UCLA, Los Angeles, CA, 90095

## Abstract

Epstein-Barr Virus (EBV) is a ubiquitous herpesvirus that typically causes asymptomatic infection but can promote B lymphoid tumors in the immune-suppressed. *In vitro*, EBV infection of primary B cells stimulates glycolysis during immortalization into lymphoblastoid cell lines (LCLs). Lactate export during glycolysis is crucial for continued proliferation of many cancer cells-part of a phenomenon known as the “Warburg effect,” and is mediated by the monocarboxylate transporters 1 and 4 (MCT1 and MCT4). However, the role of MCT1/4 has yet to be studied in EBV-associated malignancies which display Warburg-like metabolism *in vitro*. Here, we show that EBV infection of B lymphocytes directly promotes temporal induction of MCT1 and MCT4 through the viral proteins EBNA2 and LMP1 respectively, with MCT1 being induced early after infection and MCT4 late. Remarkably, singular MCT1 inhibition early, and dual MCT1/4 inhibition in LCLs using a novel MCT4-selective inhibitor led to growth arrest and lactate buildup. Metabolic profiling in LCLs revealed significatly reduced oxygen consumption rates (OCR) and NAD+/NADH ratios, contrary to prevous observations of increased OCR and unaltered NAD+/NADH ratios in MCT1/MCT4-inhibited cancer cells. Furthermore, U-^13^C6 glucose labeling of MCT1/4-inhibited LCLs also revealed increased labeling of glutathione in the presence of elevated ROS and depleted glutathione pools, as well as increased labeling of *de novo* pyrimidine biosynthetic intermediates, suggesting broad effects on LCL metabolism. These vulnerabilities sensitized LCLs as well as EBV+, and the related gammaherpesvirus KSHV+ lymphoma cell lines to killing by metformin and phenformin, pointing at a novel therapeutic approach for viral lymphomas.

## INTRODUCTION

Epstein-Barr Virus (EBV) is a ubiquitous human herpesvirus transmitted through saliva, with more than 95% of the world being infected by adulthood. Primary EBV infection is typically asymptomatic and surmounts to latency establishment in quiescent memory B cells in the peripheral blood(1). However, in immunocompromised persons, EBV reactivation can cause uncontrolled latent EBV-induced B-cell proliferation and frank lymphoma. Recent evidence suggests that host responses such as the DNA damage response (DDR) and metabolic stress play a significant cell-intrinsic role in preventing EBV-driven B-cell transformation(2, 3).

EBV infection of primary human peripheral blood B cells *in vitro* produces indefinitely proliferating lymphoblastoid cell lines (LCLs). LCLs express the Latency III growth program, where all six EBV Nuclear Antigens, or EBNAs, (EBNA-LP, 1, 2, 3A, 3B, and 3C) and two Latent Membrane Proteins, or LMPs, (LMP1 and LMP2A/B) are present. This gene expression program mimics that of many EBV-associated B-lymphoid cancers, making LCLs a suitable model for studying mechanisms underlying tumorigenesis(4). Prior to Latency III establishment in LCLs and shortly after EBV infection of B cells, a transient period of hyperproliferation, called Latency IIb, is induced, where only the viral EBNAs are expressed in the absence of the LMPs(5, 6). This culminates in an irreversible growth arrest in most cells caused by a persistent DNA Damage Response (DDR)(7). However, a very small subset (<10%) of hyperproliferating cells evade host barriers like the DDR, and transition into Latency III-expressing LCLs within weeks(2).

During EBV immortalization of B cells into LCLs, increased energetic demands for continued proliferation drive an upregulation of glycolysis and extracellular acidification rates (ECAR)(3, 8–10). While the ECAR is largely a reflection of lactate export into the extracellular environment, the magnitude and role of lactate export during EBV-driven B-cell immortalization remains poorly characterized. Besides being a glycolytic waste product, lactate has recently emerged to play a key role in tumor development, growth, and metastasis(11, 12). Furthermore, intracellular lactate buildup can have dire consequences on cell homeostasis due to acidification by lactic acid, highlighting the importance of lactate export for sustained cell growth and survival(13).

Proton-coupled lactate export is driven by the plasma membrane proteins monocarboxylate transporters 1 and 4 (MCT1 and MCT4). MCT1 has a much higher affinity for lactate than MCT4, making it the primary lactate transporter under normal physiological conditions (14–16). MCT4, however, is typically expressed in highly glycolytic tissues, or when intracellular lactate concentrations are high, such as during aerobic glycolysis(17–19). MCT1 and MCT4 have the ability to transport other monocarboxylates such as pyruvate and ketone bodies like acetoacetate and β-hydroxybutyrate(15, 19, 20). However, these transporters are most commonly associated with their role in lactate transport across the plasma membrane, especially during tumor development, growth, and metastasis(21–23). As such, MCT1 and MCT4 act as essential buffers at the intersection of the intracellular and extracellular environments, sensing monocarboxylate concentrations and facilitating rapid transport based on the needs of the cell(24–26). Not surprisingly, MCT1 and MCT4 are commonly expressed in a wide variety of tumors, exporting lactate from glycolytic cancer cells, which can later be recycled by oxidative cells in the tumor microenvironment(27, 28).

Targeting MCT1 and MCT4-mediated lactate export has become a highly attractive therapeutic strategy in recent years, as blocking lactate export can lead to cell cycle arrest, and sensitize tumor cells to killing by other metabolic inhibitors(20, 22, 29, 30). MCT1 inhibitors are currently in Phase I clinical trials for a number of solid tumors and lymphomas(31). While these inhibitors have proven to effectively reduce tumor burden in preclinical models of MCT1-overexpressing cancers, they are unfortunately ineffective in MCT1/MCT4-co-expressing cancers due to the redundant role of MCT4 as a lactate exporter(32). This highlights the need for MCT4-specific inhibitors, which are unfortunately not commercially available to date. Furthermore, while inhibition of MCT1 and MCT4 universally leads to a buildup of intracellular lactate in tumor cells, the mechanisms underlying growth arrest and cell death are not fully understood. Thus, studying how MCT1 and MCT4 are regulated in diverse contexts is critical for expanding our understanding of their role not only in tumorigenesis, but cell biology and homeostasis at large.

Given their established roles in tumor development both *in vitro* and *in vivo*, we reasoned that MCT1 and MCT4-mediated lactate export might contribute to EBV-mediated B-cell tumorigenesis, and virus-associated lymphoma cell growth more broadly. To address this, we measured changes in lactate concentration and MCT expression through EBV-mediated B-cell outgrowth and perturbed MCT1 and MCT4 function with small molecule antagonists. Our data identify metabolic vulnerabilities that we propose could be exploited for the treatment of viral-associated cancers.

## RESULTS

### EBV infection of human B cells leads to an increase in lactate accumulation, secretion, and the lactate transporter MCT1

We and others have previously demonstrated that EBV infection of primary human B cells leads to upregulation of glycolysis as evidenced by an increase in glycolytic enzymes and the extracellular acidification rate (ECAR)(3, 10). To determine whether the increase in ECAR was indicative of an increase in lactate export from infected cells, we harvested media from EBV-infected primary human B cells at several time points following infection and measured media lactate. We found that media lactate levels increased steadily after infection as cell proliferation initiated by day 4 and through long-term outgrowth (day 35 = lymphoblastoid cell line, or LCL) (**Fig. 1A**). However, we did not observe a concomitant rise in intracellular lactate until late times post infection (**Fig. 1B**). This finding led us to assess the mechanism of lactate secretion during early infection.

**Figure 1.**
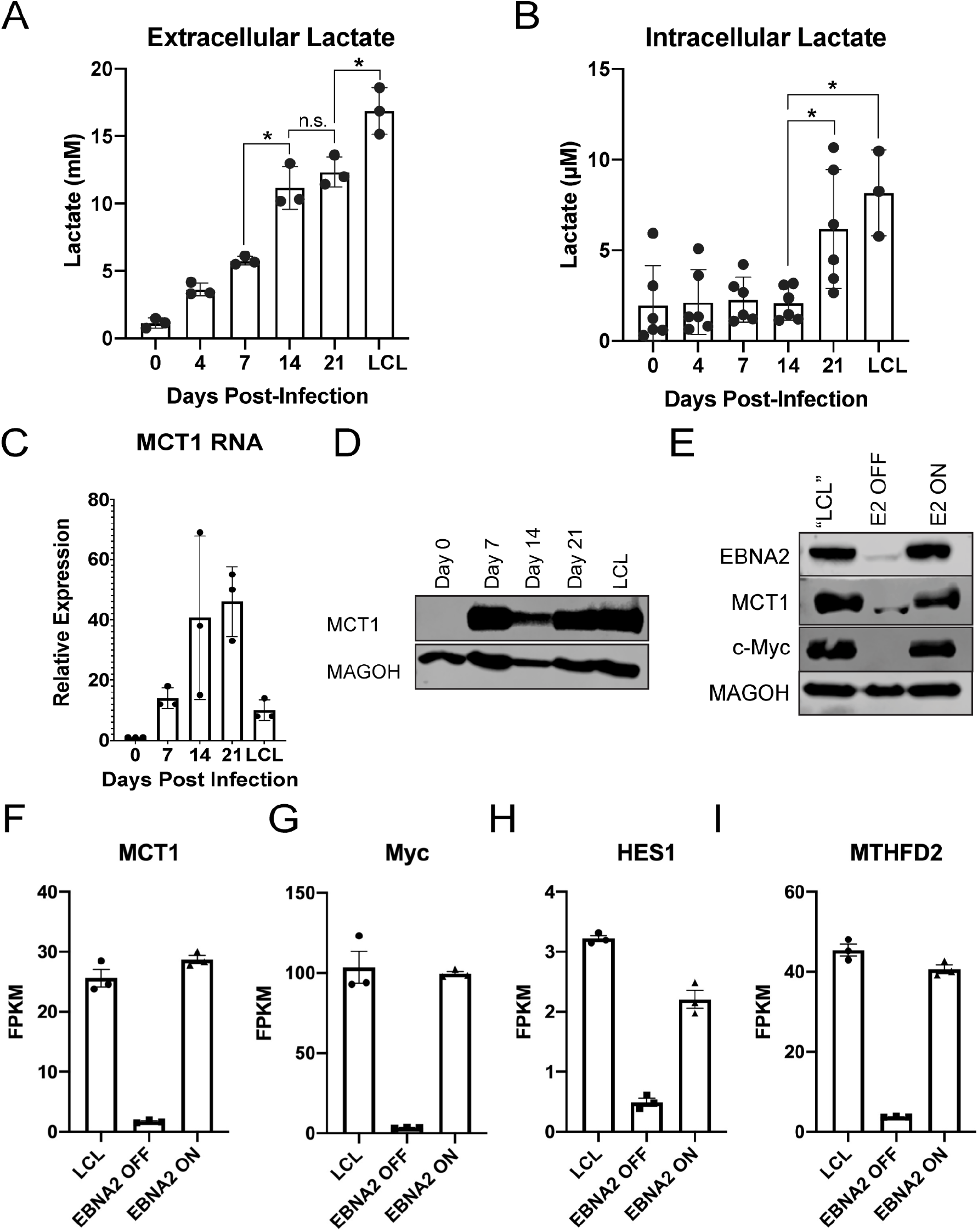
EBV infection of human B cells leads to an increase in lactate transport through MCT1. ***A***. Extracellular (n=3) and ***B***. Intracellular (n=6) lactate concentration during EBV-infected B-cell immortalization. Extracellular concentration was obtained from the supernatant of CD19+ isolated B cells at a concentration of 1×10^6^/mL and corrected to cell-free growth medium lactate concentrations. Intracellular lactate concentration is per 12,500 cells. ***C***. MCT1 RNA (n=3) and ***D***. Representative protein levels during EBV-infected B-cell immortalization. MAGOH is a loading control used because the expression of this gene is not changed upon B-cell outgrowth. ***E***. Immunoblot of EBNA2, MCT1, and c-Myc in P493-6 cells at the “LCL” state with stable EBNA2 expression, EBNA2 OFF in the absence of β-estradiol, and EBNA2 ON with EBNA2 expression regained. ***F-I***. RNA-Seq data showing MCT1, c-Myc. HES1, and MTHFD2 expression in P493-6 B cells (n=3). FPKM= Fragments Per Kilobase of transcript per Million mapped reads. Statistical significance determined by paired t-test. *=p<0.05, **=p<0.01, ***=p<0.001.

Under normal physiological conditions in most cells, lactate transport is mediated by the monocarboxylate transporter 1 (MCT1). We found that while resting B cells had little MCT1 expressed at the RNA or protein level, by day 7 EBV-infected B cells were expressing high levels of MCT1 that persisted through LCL outgrowth (**Fig. 1C-D**). MCT1 is a known transcriptional target of c-Myc, which is transcriptionally regulated upon EBV infection by the latency protein EBV Nuclear Antigen 2 (EBNA2)(33). To test the hypothesis that EBNA2 is the viral protein responsible for MCT1 induction, we utilized the P493-6 model of regulated EBNA2 expression(34). In this system, an endogenous EBNA2 is regulated post-translationally by β-estradiol due to its fusion to the estrogen receptor. Loss of EBNA2 in this system in the absence of β-estradiol led to a loss of MCT1 as well as c-Myc mRNA and protein and the mRNA of two other EBNA2 and c-Myc targets, HES1 and MTHFD2 (**Fig. 1E-I**) (10, 35). Similarly, turning EBNA2 back on in these cells led to re-expression of these targets.

### MCT1 inhibition causes growth arrest and increased lactate in early EBV-infected B cells

The induction of MCT1 expression early after EBV infection led us to ask whether MCT1 was crucial for maintaining EBV-infected B-cell proliferation or survival given the known detrimental consequences of lactate accumulation(13). To assess this, we used the MCT1 inhibitor, AZD3965, which is currently in clinical trials for MCT1-overexpressing lymphomas(31). We treated EBV-infected peripheral blood mononuclear cells (PBMCs) concurrent with infection and assessed CD19+ B-cell proliferation by flow cytometry using the proliferation tracking dye CellTrace Violet at different days post infection. We found that treatment with AZD3965 led to a dose-dependent decrease in B-cell proliferation on days 4 and 7 post EBV infection, accompanied by an increase in intracellular lactate levels (**Fig. 2A-C**). This effect on proliferation was due to a G1/S phase growth arrest and not to increased B-cell death (**Supp. 1**). However, this effect was lost by 14 days post infection and later in immortalized LCLs, suggesting an acquired resistance to MCT1 inhibition (**Fig. 2D-E**).

**Figure 2.**
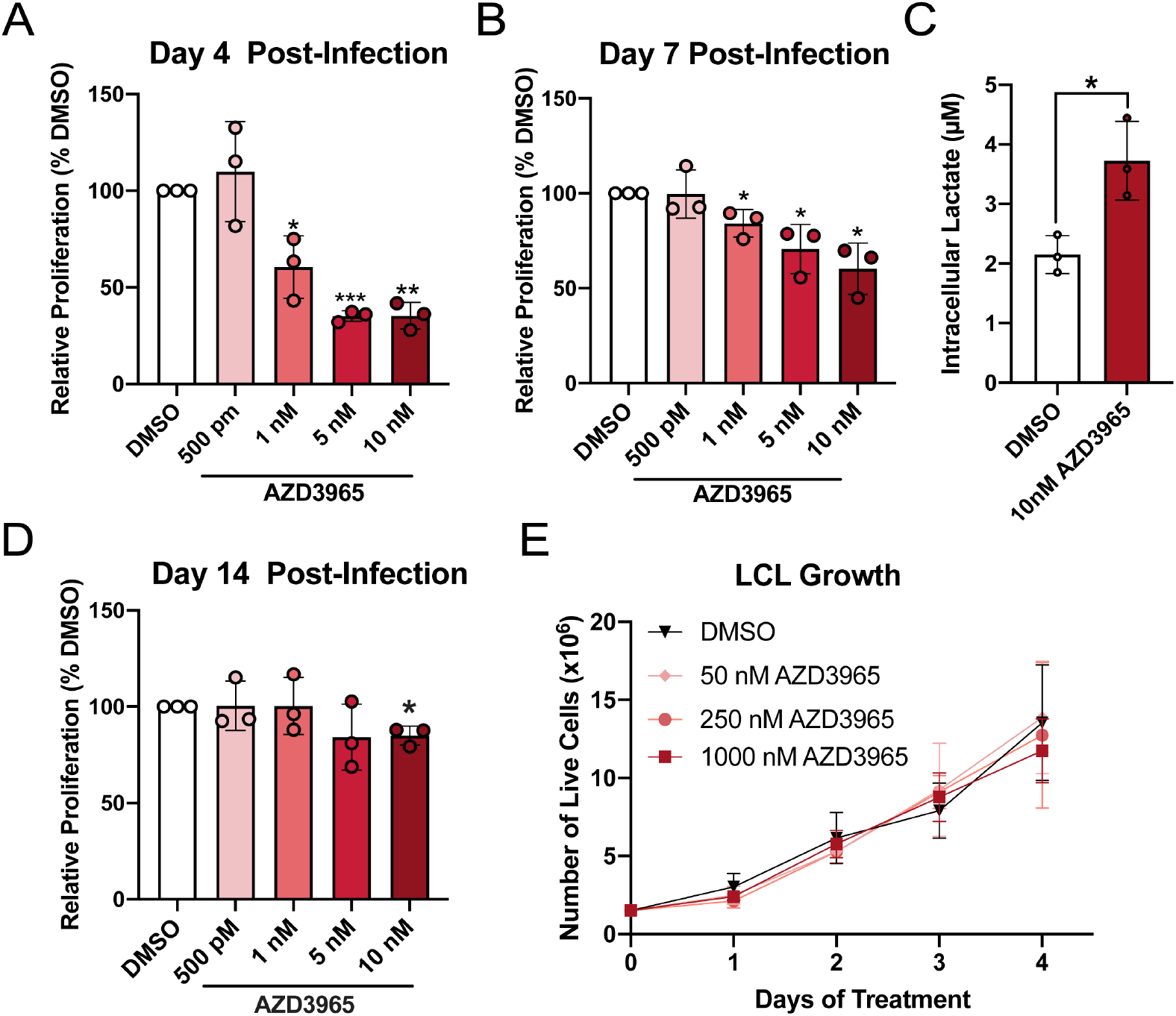
MCT1 inhibition leads to growth arrest and lactate buildup in early EBV-infected B cells. ***A***. Bulk PBMCs (n=3) were treated with AZD3965 immediately following EBV infection, and CD19+ B-cell proliferation was determined by flow cytometry 4 days later. Cell counts were bead normalized. ***B***. Bulk PBMCs (n=3) were treated 4 days post-infection, and CD19+ B-cell proliferation assessed by flow cytometry at 7 days post-infection. ***C***. CD19-purified B cells (n=3) were treated at four days post-infection and intracellular lactate assessed three days later at seven days post-infection. ***D***. Same as ***B***, except proliferation was assessed at 14 days post-infection. ***E***. Growth curve of LCLs (n=3) treated with varying concentrations of AZD3965. LCL growth was determined by daily assessment of trypan blue exclusion. Statistical significance determined by paired t-test. *=p<0.05, **=p<0.01, ***=p<0.001.

### EBV-infected B cells acquire resistance to MCT1 inhibition during B-cell transformation through induction of MCT4

Given the fact that our data indicate a significant increase in intracellular lactate concentrations in LCLs compared to early-infected B cells, we hypothesized that EBV immortalization might induce the expression of another lactate exporter, which would mitigate the effects of MCT1 inhibition. We found that expression of the MCT1 family member, MCT4, significantly increased between days 7 and day 14 post-EBV infection (**Fig. 3A-C**). Interestingly, this time period correlates with a known switch in the EBV latency gene expression program from latency IIb (EBNAs only - EBNA1, 2, 3A, 3B, 3C, and LP) to latency III (EBNAs and LMPs – LMP1, LMP2A/B). Indeed, LMP1 expression within LCLs correlated with MCT4, but not MCT1 or MCT2 expression (**Fig. 3D**). Furthermore, MCT4 was found to be upregulated by LMP1 in a publicly available RNA-Seq dataset of LMP1-expressing DG75, EBV-negative Burkitt lymphoma cells (**Fig. 3E**). Thus, the switch to latency III and expression of the viral protein LMP1 leads to B-cell induction of MCT4 that may help compensate for elevated glycolysis in EBV-infected cells as they progress to become LCLs.

**Figure 3.**
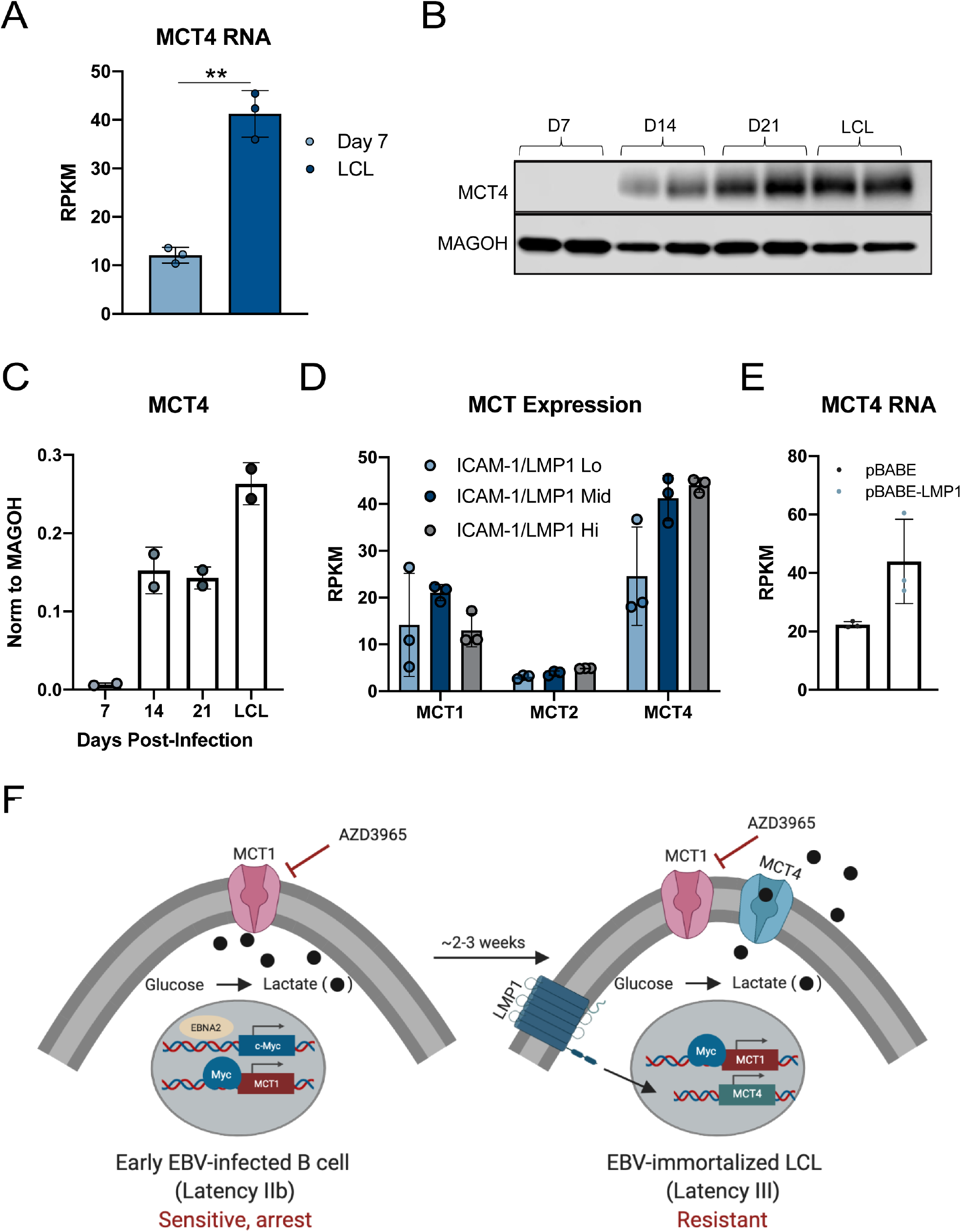
EBV-infected B cells acquire resistance to MCT1 inhibition during B-cell transformation due to MCT4 induction. ***A***. Queried RNA-Seq data of MCT4 expression in EBV-infected B cells (n=3) at 7 days post-infection, and LCL(62). RPKM= Reads Per Kilobase of transcript per Million mapped reads. ***B***. Western blot of MCT4 in CD19+-purified B cells during EBV-mediated B-cell immortalization. MAGOH= loading control. ***C***. quantification of B (n=2). ***D***. Monocarboxylate transporter (MCT) expression from RNA-Seq of LCLs sorted by the LMP1 proxy, ICAM-1, status. ***E***. Queried RNA-Seq data of MCT4 expression in DG75 cells (n=3) transfected with LMP1. Statistical significance was determined by paired t-test. *=p<0.05, **=p<0.01, ***=p<0.001. ***F***. Schematic representation of MCT1 inhibitor sensitivity during EBV-mediated B lymphocyte immortalization.

### Dual MCT1/4 inhibition suppresses LCL growth and sensitizes to killing by ETC inhibitors

The unique expression profile of MCT1 with MCT4 at later stages of EBV-mediated B-cell outgrowth along with the significant accumulation of lactate in LCLs, led us to hypothesize that MCT4 might play a critical role in LCL growth. To assess the role of MCT4 in LCLs, we took advantage of a novel, low-nanomolar MCT4 inhibitor, VB124, which displays >1,000-fold selectivity over MCT1(36). Much to our surprise, MCT4 inhibition had no observed effect on LCL growth or viability (**Fig. 4A**). However, we reasoned that though MCT4 expression was unique to LCLs during EBV-mediated B-cell outgrowth, perhaps the presence of MCT1 in LCLs rendered them resistant to singular MCT4 inhibition. To test this, we treated LCLs with both AZD3965 (1 μM) and VB124 (20 μM) and monitored cell proliferation. Dual inhibition of MCT1 and MCT4 led to a significant decrease in LCL growth, accompanied by a four-fold increase in intracellular lactate (**Fig. 4B, C**). Similar to the MCT1 inhibition effect on early-infected B-cell proliferation, MCT1/4 combined inhibition led to growth arrest in LCLs rather than cell death (**Supp. Fig 2**). Treatment with exogenous L-lactate phenocopied dual MCT1/4 inhibition suggests a direct role for lactate accumulation in the regulation of EBV-immortalized LCL growth (**Supp. Fig 3**).

**Figure 4.**
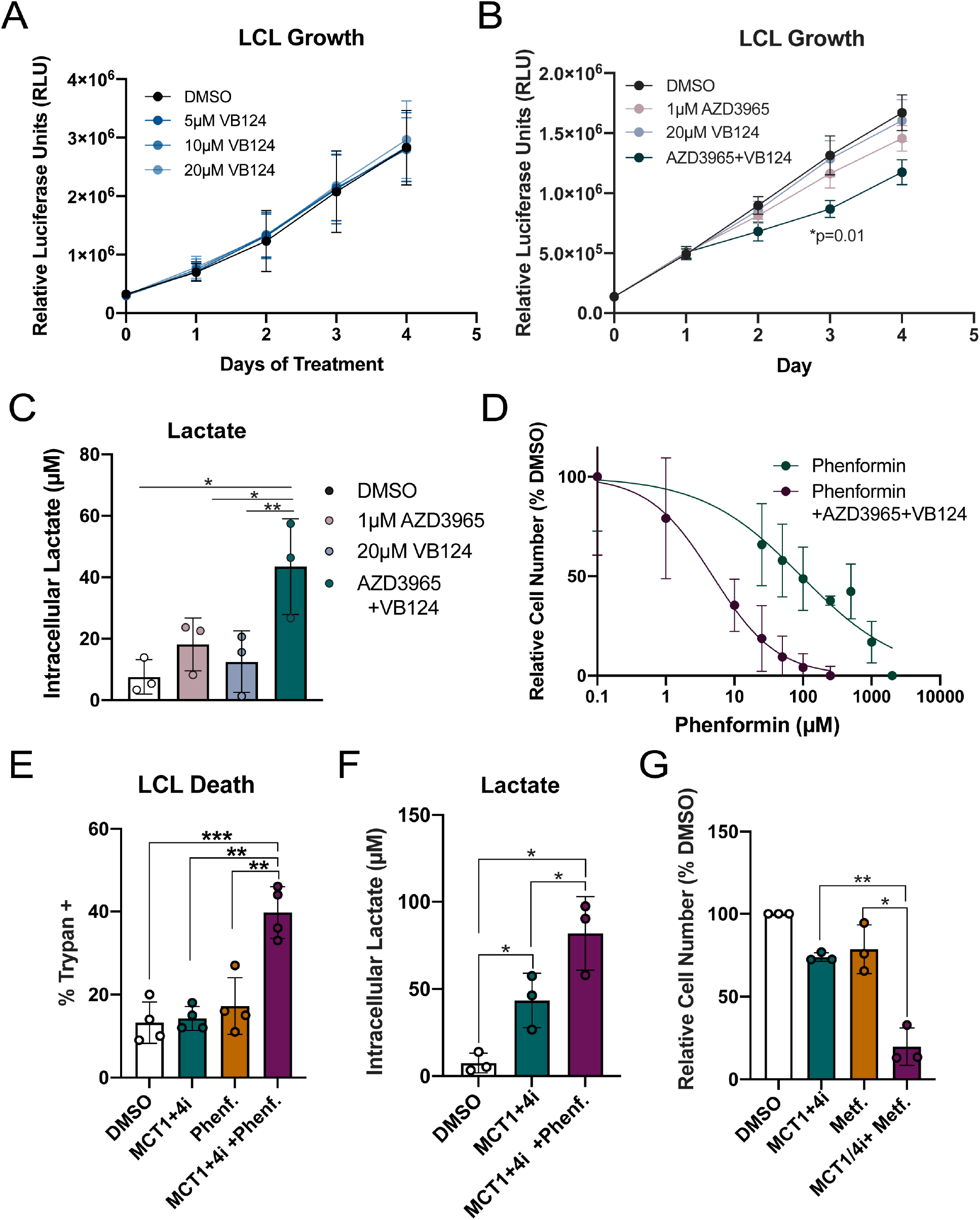
Dual MCT1/4 inhibition sensitizes LCLs to killing by electron transport chain (ETC) inhibitors. ***A***. Growth curve of LCLs (n=3) treated with varying concentrations of VB124. Growth was determined by CellTiter Glo. ***B***. Growth curve of LCLs (n=3) treated with either DMSO, 1μM AZD3965, 20μM VB124, or 1μM AZD3965 + 20μM VB124. Growth was assessed as in A. ***C***. Intracellular lactate concentration (per 12,500 cells), of LCLs (n=3) treated with either DMSO, 1μM AZD3965, 20μM VB124, or 1μM AZD3965 + 20μM VB124. ***D***. Phenformin IC50 curve. Generated 72h after treatment of LCLs (n=3) with either phenformin or 1μM AZD3965 + 20μM VB124. Relative cell number was determined by CellTiter Glo luminescence. Values were normalized to DMSO to obtain %DMSO. ***E***. LCL death, as determined by trypan blue staining after 72h of treatment. n=3. MCT1+4i= 1μM AZD3965 + 20μM VB124, Phenf. = 10μM Phenformin ***F***. Intracellular lactate concentration (per 12,500 cells), of LCLs treated for 3 days ***G***. Relative LCL number as determined by CellTiter Glo, 72h after treatment. n=3. MCT1 +4i= 1μM AZD3965 + 20μM VB124, Metf. = 2mM Metformin. Statistical significance was determined by paired t-test. ***H***. LCL death, based on trypan blue positivity, following 72h of treatment. n=3. *=p<0.05, **=p<0.01, ***=p<0.001.

We next sought to assess whether MCT inhibition might also render LCLs susceptible to arrest or killing by electron transport chain (ETC) inhibitors as this therapeutic strategy has been successful in other cancer models(30, 37). Indeed, LCLs were 20-fold more susceptible to the complex I inhibitor phenformin (IC50 from ~100μM to ~5μM) when coupled with growth inhibitory concentrations of AZD3965 (1μM) and VB124 (20 μM) (**Fig. 4D**). Moreover, treatment of LCLs with 10 μM phenformin induced significantly more cell death in MCT1/4-inhibited cells than alone (**Fig. 4E**). This increased cell death was accompanied by increased intracellular lactate levels as well (**Fig. 4F**). Finally, we found that MCT1/4 inhibition also sensitized LCLs to killing by the FDA-approved ETC inhibitor, metformin (**Fig. 4G-H**). These experiments support our hypothesis that lactate accumulation in EBV-infected cells renders them hyper-sensitive to the ETC inhibitors phenformin and metformin.

### Dual inhibition of MCT1/4 in LCLs compromises OCR and NAD^+^/NADH ratios, and induces oxidative stress

We next sought to explore the mechanisms responsible for the sensitivity of LCLs to MCT inhibition. We performed Seahorse assays on LCLs treated with MCT1 and 4 inhibitors for 24 hours. As expected, dual MCT1/MCT4 inhibition, which induced lactate accumulation in LCLs, significantly decreased extracellular acidification (**Fig. 5A**). Unexpectedly, however, dual MCT1/4 inhibition significantly decreased oxygen consumption rates (OCR) after just 24 hours of treatment indicating that these metabolic changes precede the growth arrest observed at later time points (**Fig. 5B**). This effect was even more pronounced with the addition of phenformin, which was expected given the unequivocal role of the ETC on mitochondrial respiration (**Fig. 5B**).

**Figure 5.**
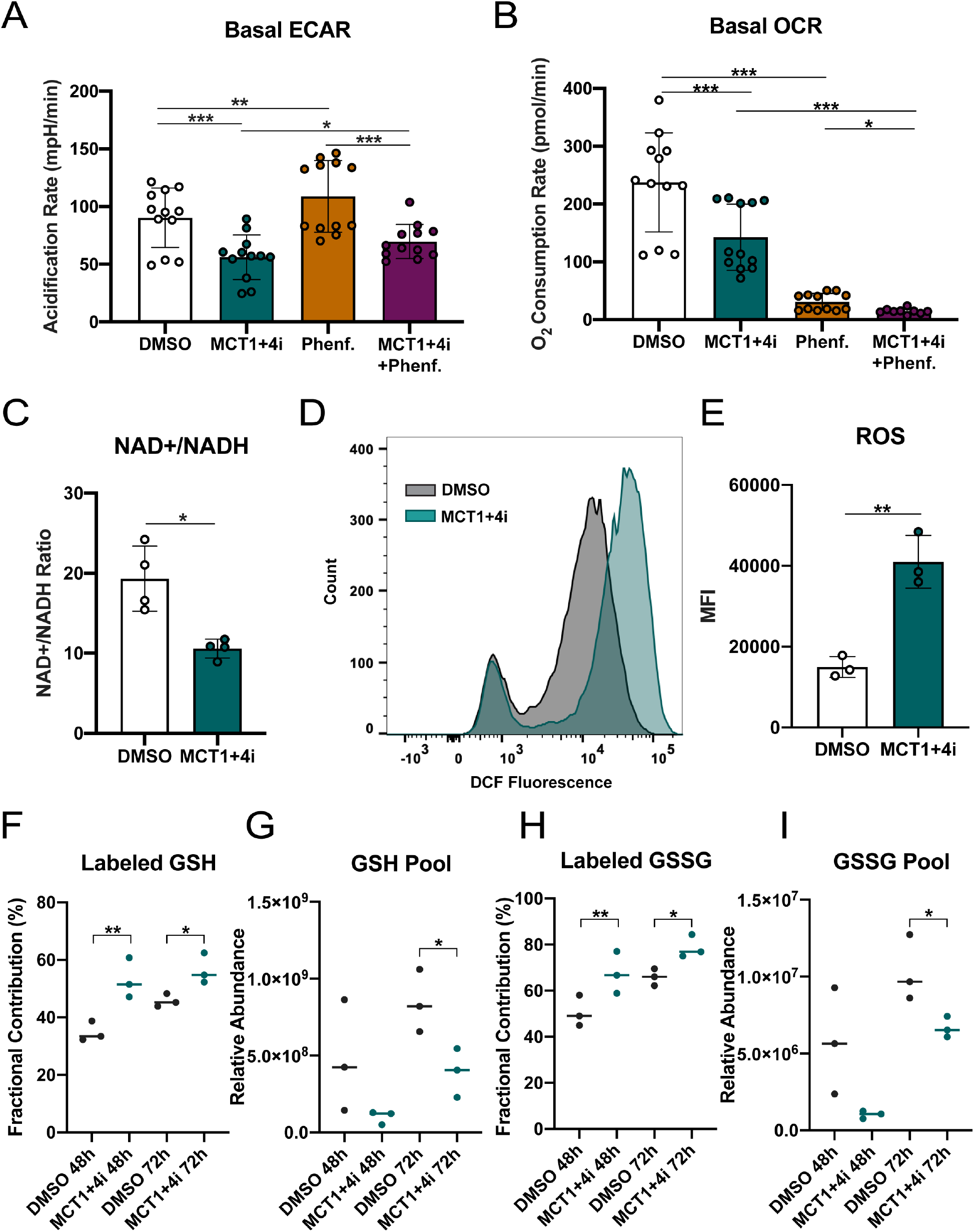
Dual inhibition on MCT1/4 in LCLs compromises OCR and NAD/NADH ratios and induces oxidative stress. ***A***. Seahorse assay showing extracellular acidification rate (ECAR) and ***B***. Oxygen consumption rate (OCR) of LCLs (n=4) in triplicate. MCT1+4i= 1μM AZD3965 + 20μM VB124. Phenf. = 10μM Phenformin. ***C***. NAD^+^/NADH ratio of LCLs (n=4) treated for 48h ***D***. Fluorescence-activated cell sorting (FACS) histograms of treated LCLs (n=3) stained with 2’,7’-dichlorodihydrofluorescein diacetate (H2-DCFDA). ***E***. Mean fluorescence intensity (MFI) of D. ***F-I***. Metabolic analysis with ^13^C-6-glucose tracing of reduced (GSH) or oxidized (GSSG) glutathione in LCLs (n=3) treated with either DMSO or MCT1+MCT4 inhibitors for 48h or 72h. Cells were cultured in the presence of ^13^C6-labeled glucose-supplemented RPMI medium. Fractional contribution= % heavy-labeled GSH or GSSG relative to pool size. Pool= Relative abundance of GSH or GSSH in sample, as determined by LC-MS. Statistical significance was determined by paired t-test. *=p<0.05, **=p<0.01, ***=p<0.001.

The production of lactate during glycolysis is coupled with the regeneration of NAD^+^ from NADH. Therefore, we assessed the NAD^+^/NADH ratio in MCT1/4 inhibited LCLs. We found that treated cells had a significantly lower NAD^+^/NADH ratio than control LCLs (**Fig. 5C**). Since NAD^+^/NADH ratios directly contribute to the redox state of the cell, we investigated whether compromised NAD^+^/NADH levels in MCT1/4-inhibited LCLs could be inducing oxidative stress. Using the reactive oxygen species (ROS)-sensitive fluorescent DCFDA dye, we found that ROS levels increased significantly upon MCT1/4 inhibition (**Fig. 5D**). Furthermore, metabolite tracing experiments using heavy-labeled ^13^C-glucose indicated that dual MCT1/4 inhibition increased the fractional contribution of glucose in reduced and oxidized glutathione (**Fig. 5F,H**). Furthermore, the marked decrease in glutathione pools by 72 hours post-treatment suggests an effort to mitigate accumulating ROS levels through glutathione antioxidant activity (**Fig. 5G,I**)

### EBV+ and KSHV+ lymphoma cell lines are sensitive to dual MCT1/4 and ETC inhibitors

EBV is an etiologic factor in the development of B-cell lymphomas of the immune suppressed such as HIV-associated diffuse large B-cell lymphomas (DLBCL) and post-transplant lymphoproliferative disease (PTLD)(38–40). Therefore, we asked whether the novel approach of dual MCT1/4 inhibition in EBV-immortalized LCLs could also be effective in EBV-positive lymphoma cells. We treated the EBV+ AIDS-immunoblastic lymphoma cell line, IBL-1, which like LCLs, expresses MCT1 and MCT4, with both MCT1/4 inhibitors (**Fig. 6C**). We observed a striking diminution in growth, similar to the phenotype in LCLs (**Fig. 6A**). Consistently, we also observed synergy between MCT1/4 antagonism and metformin as well as phenformin in IBL-1 cells, which supports a novel therapeutic approach to EBV+ lymphomas (**Fig. 6A and 6B**).

**Figure 6.**
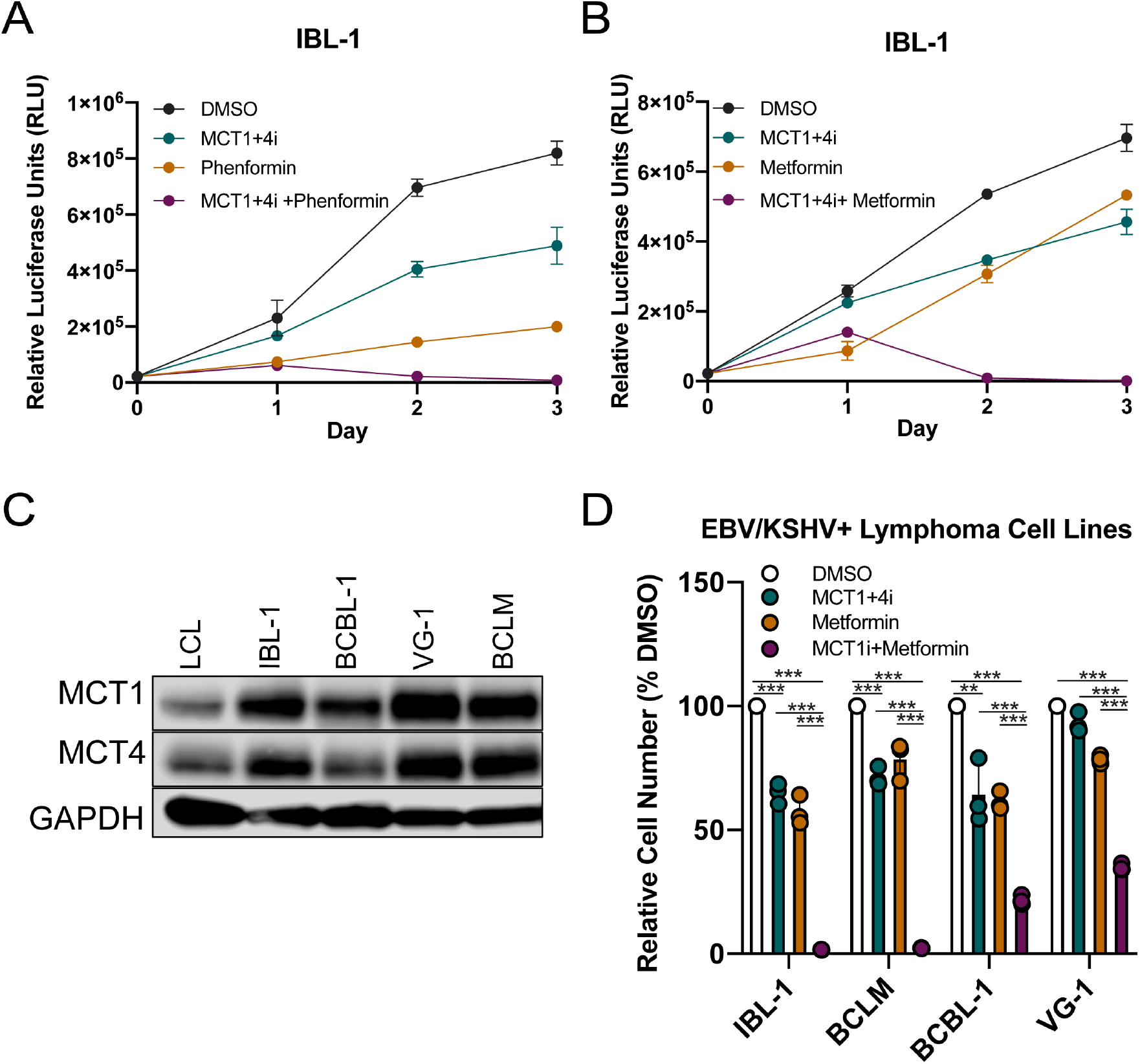
EBV+ and KSHV+ lymphoma cell lines are sensitized to killing by metformin upon dual MCT1/4 inhibition. ***A, B***. Growth over time of the EBV+ AIDS immunoblastic lymphoma cell line IBL-1 (n=3), in the presence of both MCT1/4 inhibitors, and ETC inhibitors metformin and phenformin. MCT1+4i= 1μM AZD3965 + 20μM VB124. Phenformin= 100μM Metformin= 2mM. CellTiter Glo luminescence was used as proxy for cell growth. ***C***. Immunoblot of MCT1 and MCT4 in EBV+ or KSHV+ primary effusion lymphoma (PEL) cell lines, compared to EBV-immortalized LCLs. IBL-1= EBV+, KSHV-AIDS immunoblastic lymphoma, BCBL-1 = EBV-, KSHV+ PEL, VG-1 = EBV-, KSHV+ PEL, BCLM= EBV-, KSHV+ PEL. GAPDH= loading control. ***D***. Relative cell count of EBV+/KSHV+ lymphoma cell lines (n=3) 48h after treatment. Statistical significance was determined by multiple t-test. *=p<0.05, **=p<0.01, ***=p<0.001.

Given the robustness of this strategy, we asked whether other viral lymphomas may be susceptible to MCT antagonism and combination with metformin. Kaposi Sarcoma Herpesvirus (KSHV) is a closely related oncogenic gamma herpesvirus to EBV and is the etiologic agent of Kaposi Sarcoma as well as primary effusion lymphomas (PEL) in HIV-infected individuals. First, we confirmed expression of MCT1 and MCT4 in the PEL cell lines BCBL-1, VG-1, and BCLM, alongside IBL-1 and LCLs (**Fig. 6C**). Then, we investigated whether dual MCT1/4 inhibition could impact PEL growth, especially when combined with metformin, as was true in LCLs and IBL-1 cells. Consistent with our observation in EBV-infected cells, we observed that dual inhibition of MCT1 and 4 uniformly suppressed growth of KSHV-infected PEL cells. Furthermore, MCT1/4 antagonism also increased sensitivity of PEL cells to metformin-mediated growth inhibition, thereby expanding the applicability of this strategy to KSHV+ lymphomas (**Fig. 6D**).

## DISCUSSION

Epstein-Barr virus reorganizes B-cell metabolic pathways to promote immortalization as a precursor to the development of lymphomas in the immune suppressed. We and others have previously described a balanced upregulation of oxidative phosphorylation and glycolysis through EBV-mediated B-cell outgrowth(3, 10). In order to maintain glycolytic flux during aerobic glycolysis, EBV-infected B cells must upregulate lactate transporters. In this study, we identified an EBNA2/c-Myc mediated upregulation of MCT1 that was important for supporting increased glycolysis during the hyper-proliferative burst early after EBV infection. However, infected cells became resistant to MCT1 antagonism between one- and two-weeks post infection and we observed a strong upregulation of MCT4. This correlated with the switch in viral latency programs in which the viral LMP1 protein promoted MCT4 expression. EBV-immortalized lymphoblastoid cell lines were sensitive to the inhibition of both MCT1 and MCT4 with specific small molecule antagonists, which led to their growth arrest due to a loss in the oxygen consumption rate, an accumulation of both lactate and reactive oxygen species, and depletion of NAD^+^ relative to NADH. Importantly, MCT1/4 inhibition rendered LCLs vulnerable to the inhibition of the electron transport chain with phenformin or metformin. This synergistic effect on LCLs was also manifested in lymphoma cell lines harboring EBV and also a related oncogenic gammaherpesvirus, KSHV, suggesting a novel strategy for targeting viral-induced cancers.

The role of monocarboxylate transporters (MCTs) has been well studied in cancer, but not during viral infection. The metabolic requirements of both processes are similar with a goal of ramping up anabolic processes to either promote proliferation or viral particle production. Latent oncogenic herpesviruses are unique in that they express specific proteins and non-coding RNAs that reprogram cellular transcription and metabolic pathways, with the goal of promoting viral genome replication during host cell proliferation. For example, during KSHV lytic replication, glycolysis is required for maximal virion production, while during latent infection, 10 out of the 12 microRNAs are sufficient to induce glycolysis, thereby contributing to infected cell growth (41,42). KSHV-associated PEL also display elevated aerobic glycolysis and fatty acid biosynthesis compared to primary B cells, highlighting the importance of these pathways in KSHV-associated tumors, and another parallel between KSHV infection and tumorigenesis(43)’(44). Similarly, the EBV oncoprotein LMP1 upregulates glucose import through NFκB signaling to promote cell survival(8). Thus, EBV and KSHV coordinately activate their infected cells in a similar manner to how cellular oncogenes cooperate to reprogram metabolism in cancer. While both EBV and KSHV-associated lymphomas display increased extracellular acidification rates (ECAR) and lactate export due to increased glycolysis, only a handful of studies have addressed lactate transporters in these cancers(9, 45). In one study, MCT4 RNA was shown to be upregulated in EBV-immortalized LCLs relative to mitogen-activated B cells and freshly EBV-infected B cells, but mainly due to its association with the induction of hypoxia(9, 46). More indirectly, KSHV-infected PEL chemoresistance to doxorubicin and paclitaxel has been linked to expression of the MCT1/4-associated chaperone glycoprotein Emmprin, which is regulated by the KSHV-encoded latency-associated nuclear antigen (LANA) in KSHV+ PEL cell lines (47)’(48). Given the similarities between EBV and KSHV-mediated reprogramming of host metabolism, and based on our observation that KSHV+ PELs arrest upon treatment with both MCT1/4 inhibitors, it is therefore possible that both viruses require and/or regulate MCT1/4-mediated lactate export in similar ways. Future studies would need to elucidate whether MCT1/4-mediated lactate transport in KSHV+ lymphomas is intertwined with other aspects of metabolism like redox regulation and oxidative stress mitigation, as is the case in EBV-immortalized LCLs.

During EBV infection, the EBV master transcriptional regulator EBNA2 can both directly and indirectly influence host metabolic pathways such as one-carbon metabolism and fatty acid biosynthesis through c-Myc(10, 49). We find that concomitant induction of MCT1 through c-Myc is critical to support continued proliferation as lactate is generated through glycolysis. Interestingly, we found that early-infected cells were exquisitely sensitive to MCT1 inhibition through AZD3965 with low nanomolar (~1-10 nM) concentrations halting growth while in many other tumor cell systems 250 nM is typically used. This suggests that MCT1-mediated lactate export could be particularly important for B-cell growth in the absence of MCT4 – a gene expression signature which has also been observed in germinal center B cells(32).

We were intrigued by the finding that EBV-infected B cells became resistant to MCT1 inhibition between one and two weeks after infection as this is the timeframe when infected cells switch from proliferation driven by the viral EBNA proteins (latency IIb) to that of the EBNAs and the LMPs (latency III)(5, 6). Given this relationship and the prior knowledge that MCT4 induction could promote resistance to MCT1 antagonism, we interrogated MCT4 upregulation and sensitivity to a novel MCT4 inhibitor. Our data support the upregulation of MCT4 by LMP1, which is also supported by a published data set of LMP1 expression in DG75 cells(50). Moreover, that study found that LMP1 targets dependent on the metabolic and DNA damage regulator, PARP1, were largely enriched in HIF1α targets. In fact, their data using the PARP1 inhibitor olaparib suggests that LMP1 induction of MCT4 was PARP1-dependent. These data also nicely dovetail with that of a previous study supporting the stabilization of HIF1α and its targets in EBV-infected cells supported a “pseudo-hypoxia” that appears to be induced by EBV(9).

The mechanism of growth arrest mediated by MCT1/4 antagonism involved a reduction in the extracellular acidification rate due to the accumulation of lactate within cells, but surprisingly also, a significant reduction in the oxygen consumption rate (OCR). Moreover, these effects were observed within 24 hours following treatment, preceding cellular growth arrest which occurs between 48-72 hours after treatment. We were intrigued by the significant decrease in OCR upon dual MCT1/4 inhibition, given that previous reports describe increased OCR upon lactate export inhibition as a means of compensating for reduced glycolysis with increased OXPHOS (20, 22)’(51, 52). One possible explantion for this observed decrease in OCR could be due to the altered redox balance via depleted NAD+/NADH, which could inhibiting NADH:ubiquinone oxidoreductase activity in complex I of the ETC(53). Another possibility might be due to the observed defect in *de novo* pyrimidine metabolism, as dihydroorotate dehydrogenase (DHODH) function is directly linked to complex III activity(54). Either way, the reduction of OCR upon selective MCT1/4 inhibition in LCLs points at a unique vulnerability that can be further exploited for therapeutic benefit in EBV-driven lymphomas in combination with ETC inhibitors like metformin. Recently, dual MCT1/4 inhibition using the anti-hypertensive drug syrosingopine was found to be synthetically lethal with metformin in leukemia and breast cancer cell lines due to depleted NAD^+^/NADH ratios(30). Interestingly, we found that in LCLs, selective inhibition of both MCT1 and MCT4 with AZD3965 and VB124 was sufficient to significantly deplete NAD^+^/NADH ratios, while addition of the metformin analog phenformin did not further deplete these ratios (not shown). This therefore suggests an alternative mechanism of metformin/phenformin-induced killing in MCT1/4-inhibited cancer cells, not directly related to NAD+/NADH metabolism.

The MCT1 inhibitor AZD3965 is currently in clinical trials for various lymphomas and solid tumors(31). However, no such trials exist for MCT4, primarily due to the lack of MCT4-specific inhibitors. While pan-MCT1/4 inhibitors do exist, these also target other metabolic pathways such as the glycolytic enzyme enolase, and mitochondrial pyruvate transport in addition to MCT1 and MCT4, which can produce adverse side effects(55, 56). Using the novel MCT4-specific inhibitor VB124 as a tool compound, we were able to explore the therapeutic potential of MCT1/4-targeted combination therapy in viral-associated lymhpomas. We recognize that pre-clinical testing of AZD3965 and VB124 in primary cells and cell lines may be an incomplete representation of lymphomagenesis *in vivo* as contributions from the tumor microenvironment are lacking. However, EBV has a narrow host tropism and is unable to infect mice or other common small animal models. Recent development of humanized mouse models of EBV infection will be useful in assessing how metabolic pathways like MCT1/MCT4-mediated lactate transport contribue to EBV-driven tumorigenesis *in vivo(57*).

## MATERIALS AND METHODS

### Cell lines, culture conditions, and viruses

Buffy coats were obtained from normal human donors through the Gulf Coast Regional Blood Center (Houston, TX) and peripheral blood mononuclear cells (PBMCs) were isolated by Ficoll Histopaque-1077 gradient (Sigma, H8889). B95-8 strain of Epstein-Barr virus was produced from the B95-8 Z-HT cell line as previously described (52). Virus infections were performed in bulk by adding 50 μL of filtered B95-8 supernatant to 1×10^6^ PBMCs.

All LCLs were kept in RPMI 1640 medium supplemented with 10% heat-inactivated fetal bovine serum (Corning), 2 mM L-Glutamine, 100 U/ml penicillin, 100 μg/ml streptomycin (Invitrogen), and 0.5 μg/mL Cyclosporine A (Sigma). The same conditions were used for the EBV-germinal-center-derived B-cell lymphoma line BJAB (obtained from George Mosialos, Aristotle University, Thessaloniki, Greece). Kaposi’s sarcoma-associated herpesvirus-positive (KSHV+)/EBV-primary effusion lymphoma (PEL) cell lines VG-1, BCLM, and BCBL1 were provided by Dirk Dittmer (University of North Carolina [UNC], Chapel Hill, NC) or Bryan Cullen (Duke University, Durham, NC) with permission from the original authors and kept in RPMI 1640 medium supplemented with 15% fetal bovine serum, 10 mM HEPES, 1 mM sodium pyruvate, and 0.05 mM β-mercaptoethanol (Sigma). The EBV-positive cell line derived from an AIDS immunoblastic lymphoma, IBL-1, was kindly provided by Ethel Cesarman (Weill Cornell Medical College, New York, NY) and kept in RPMI 1640 supplemented with 20% fetal bovine serum. P493-6 cells (a kind gift of Dr. Georg Bornkamm, Helmholtz Zentrum München) were cultured in RPMI 1640 supplemented with 10% tetracycline-free FBS (Hyclone SH30070), 1 μM β-Estradiol, and 1 μg/mL Tetracycline. P493-6 cells were washed three times with β-Estradiol-free media, and then cultured in the absence of β-Estradiol for 3 days to induce the EBNA2-OFF state. After washing 3 times, the cells were then cultured again for 3 days upon reintroducing 1 uM β-Estradiol, returning P493-6 cells to the EBNA2-ON state. All cells were cultured at 37°C in a humidified incubator at 5% CO2.

### Flow cytometry and sorting

To track proliferation in EBV-infected B cells, cells were stained with CellTrace Violet (Invitrogen, C34557), a fluorescent proliferation-tracking dye. Cells were first washed in FACS buffer (5% FBS in PBS), stained with the appropriate antibody for 30min-1hr at 4°C in the dark, and then washed again before being analyzed on a BD FACS Canto II. Proliferating infected B cells were sorted to a pure population of CD19+/CellTrace Violet on a MoFlo Astrios Cell Sorter at the Duke Cancer Institute Flow Cytometry Shared Resource. Mouse anti-human CD19 antibody (Biolegend Cat# 152410) conjugated with APC was used as a surface B-cell marker in flow cytometry.

### Cell Growth Assays

LCLs were seeded at a density of 3×10^5^ cells/mL, and treated with either DMSO, AZD3965 (Cayman Chemicals), VB124 (a kind gift of Mark Parnell, Vettore LLC), phenformin (Cayman Chemicals), or metformin (Cayman Chemicals) for one to four days. VB124 required concentrations much higher than its IC50 (8.6 nM) in serum due to high(~99%) serum protein binding common for carboxylic acids like VB124. Drug treatments were normalized across conditions to contain 0.01% DMSO. Cell growth was assessed daily by either CellTiter Glo (Promega) or counting by Trypan exclusion on an automated cell counter (Countess II, ThermoFisher). Luminescence was measured using a microplate reader (Biotek Synergy HTX).

### Gene Expression Analysis

Total RNA was isolated from cells by using a Qiagen RNeasy kit and then reverse transcribed to generate cDNA with the High Capacity cDNA kit (Applied Biosystems). Quantitative PCR was performed by using SYBR green (Quanta Biosciences) in an Applied Biosystems Step One Plus instrument. Primer sets include MCT1 (**F:** GACCTTGTTGGACCCCAGAG, **R**:AGCCGACCTAAAAGTGGTGG), MCT4 (**F:** AGGTCTACCTCACCACTGGG, **R**: CACAGGAAGACAGGGCTACC.

### BrdU Staining

Cell cycle progression was assessed by BrdU staining. Infected PBMCs treated with DMSO, Chk2 inhibitor II (EMD Millipore), or AZD3965 (Cayman Chemicals) cells were stained with CD19, and pulsed with BrdU (BD Biosciences, Cat# 552598) for two hours following fixation and permeabilization. Afterwards, cells were treated with a DNAse to expose BrdU-bound epitopes, and stained wth a BrdU-fluorescent antibody as well as the total DNA and cell death marker 7-AAD. Then, BrdU and 7-AAD fluorescence values were acquired on a flow cytometer (FACS Canto II).

### Immunoblotting

Cells were pelleted and washed in PBS, and then lysed in LDS Sample Buffer (NuPAGE) with complete protease inhibitors. All protein lysates were run on NuPage 4–12% gradient gels (LifeTechnology) and transferred to PVDF membrane (GE Healthcare). Membranes were blocked in 5% milk in TBST and stained with primary antibody overnight at +4°C, followed by a wash and staining with secondary HRP-conjugated antibody for 1 hour at room temperature. Antibodies include MCT1 (Abcam, Cat# ab85021, 1:1000 dilution), MCT4 (Proteintech, Cat# 22787-1-AP, 1:1000 dilution), LMP1 (S12; gift from Elliot Kieff, Harvard Medical School), c-Myc (Santa Cruz Biotechnology, SC-764), EBNA2 (PE2; gift from Elliot Kieff), MAGOH (Santa Cruz, SC-56724). We use MAGOH as a loading control because it does not change in expression from resting B cells through LCL outgrowth.

### Seahorse Analysis

Extracellular acidification rate (ECAR) and oxygen consumption rate (OCR) were measured using the Seahorse XFe96 extracellular flux analyzer (Agilent Technologies) Cell Energy Phenotype Test. LCLs were attached to culture plates by using Cell-Tak (BD Bioscience). ECAR and OCR were measured in Seahorse XF Base Medium supplemented with 1 mM pyruvate, 2 mM glutamine, and 10 mM glucose (Sigma Aldrich). ECAR and OCR values were normalized to cell number. For stress measurements, ECAR and OCR were measured over time after injection of oligomycin and FCCP.

### ^13^C-glucose Tracing

Glucose tracing experiments were performed using ^13^C6-glucose (Cambridge Isotope Laboratories Cat. CLM-1396-PK). LCLs were cultured in glucose-free RPMI supplemented with 10mM ^13^C6-glucose and 10% dialyzed FBS, seeded at a density of 300,000 LCLs/mL and treated with either DMSO or 1μM AZD3965 + 20μM VB124 for 48 or 72 hours. Then cells were harvested and washed twice with ice-cold PBS, followed by metabolite extraction with ice-cold 80% methanol in deionized water, and evaporation for approximately two hours. Dried metabolites were reconstituted in 100 μL of a 50% acetonitrile(ACN) 50% dH20 solution. Samples were vortexed and spun down for 10 min at 17,000g. 70 μL of the supernatant was then transferred to HPLC glass vials. 10 μL of these metabolite solutions were injected per analysis. Samples were run on a Vanquish (Thermo Scientific) UHPLC system with mobile phase A (20mM ammonium carbonate, pH 9.7) and mobile phase B (100% ACN) at a flow rate of 150 μL/min on a SeQuant ZIC-pHILIC Polymeric column (2.1 × 150 mm 5 μm, EMD Millipore) at 35°C. Separation was achieved with a linear gradient from 20% A to 80% A in 20 min followed by a linear gradient from 80% A to 20% A from 20 min to 20.5 min. 20% A was then held from 20.5 min to 28 min. The UHPLC was coupled to a Q-Exactive (Thermo Scientific) mass analyzer running in polarity switching mode with spray-voltage=3.2kV, sheath-gas=40, aux-gas=15, sweep-gas=1, aux-gas-temp=350°C, and capillary-temp=275°C. For both polarities mass scan settings were kept at full-scan-range=(70-1000), ms1-resolution=70,000, max-injection-time=250ms, and AGC-target=1E6. MS2 data was also collected from the top three most abundant singly-charged ions in each scan with normalized-collision-energy=35. Each of the resulting “.RAW” files was then centroided and converted into two “.mzXML” files (one for positive scans and one for negative scans) using msconvert from ProteoWizard(58). These “.mzXML” files were imported into the MZmine 2 software package(59). Ion chromatograms were generated from MS1 spectra via the built-in Automated Data Analysis Pipeline (ADAP) chromatogram module and peaks were detected via the ADAP wavelets algorithm(60). Peaks were aligned across all samples via the Random sample consensus aligner module, gap-filled, and assigned identities using an exact mass MS1(+/-15ppm) and retention time RT (+/-0.5min) search of our in-house MS1-RT database. Peak boundaries and identifications were then further refined by manual curation. Peaks were quantified by area under the curve integration and exported as CSV files. The peak areas were additionally processed via the R package AccuCor to correct for natural isotope abundance(61). Peak areas for each sample were normalized by the measured area of the internal standard trifluoromethanesulfonate (present in the extraction buffer) and by the number of cells present in the extracted sample.

### Lactate and NAD^+^/NADH Measurements

Intracellular and extracellular lactate concentrations were determined using the Lactate Glo (Promega) assay kit. Briefly, cells were harvested and washed in ice cold PBS once, then immediately inactivated with 0.6N HCl to halt metabolic activity and neutralized with 1M Tris base. The resulting lysate was combined at a 1:1 ratio with the lactate detection reagent, and luminescence measured a microplate reader. For intracellular lactate measurements involving early-infection time points, PBMCs were first depleted for CD3+ T cells (RosetteSep™ Human Granulocyte Depletion Cocktail), sorted to a pure population of CD19+ B cells on a MoFlo Astrios Cell Sorter at the Duke Cancer Institute Flow Cytometry Shared Resource. Purified B cells were either treated as indicated, or used as the starting cell population for time-course lactate measurments. Mouse anti-human CD19 antibody (Biolegend Cat# 152410) conjugated with APC was used as a surface B-cell marker in flow cytometry.

NAD^+^/NADH ratios were measured using the NAD^+^/NADH Glo assay (Promega) according to the manufacturer’s protocol. Briefly, harvested cells were washed in PBS, lysed with Tris Base supplemented with 1% Dodecyl trimethylammonium bromide, treated with 0.4N HCl (for NAD^+^), and heated (for NADH) at 60°C for 15 min. Each sample was then added at a 1:1 ratio with a luciferin detection reagent, and NAD^+^ or NADH luminescence measured separately using a BioTek Synergy™ 2 microplate reader.

### ROS Measurements

ROS fluorescence was determined using the ROS-sensitive dye DCFDA (Cayman Chemicals Cat #601520. Cells were pelleted and washed in the provided wash buffer, and then stained in 20μM DCFDA for 1.5 hours at 37°C protected from light. Then, cells were washed once in the provided wash buffer and FITC fluorescence measured in a flow cytometer. Pyocyanin (1mM) was used as a positive control and N-acetylcysteine (300mM) as a negative control.

## ACKNOWLEDGMENTS

This work was supported by National Institute of Health (NIH) grants R01-CA140337 (to M.A.L.), T32GM007105-43, T32-CA009111 (to E.N.B.), T32CA009056 (to J.C.), and R01 CA215185 (to H.R.C.) as well as RSG-16-111-01-MPC from the American Cancer Society (to H.R.C.), the CDI Harry Winston Fellowship award (to J.C.), and the St. Baldrick’s Foundation Fellowship award (to J.C.). Additional funding came from the Duke CFAR, and NIH-funded program (grant 5P30-Al064518).

The authors would like to thank Dr. Michael Cook, Nancy Martin, Lynn Martinek and the Duke Cancer Institute Shared Flow Cytometry core facility for invaluable assistance with flow cytometry and cell sorting. We would also like to thank Dr. Nancie McIver and Amanda Nichols of the Duke Cellular Metabolism Analysis Core for their guidance with the Seahorse XF anaysis. Finally, we would like to thank Ernst Schmid of the UCLA Department of Biological Chemistry for assistance with mass spectrometry and downstream analysis of isotopic tracing experiments.

## AUTHOR CONTRIBUTIONS

M.A.L., E.N.B, J.E.M., J.M.C., H.R.C., and J.C. conceived and designed research experiments. Specifically, M.A.L., J.E.M., and H.R.C. recognized that EBNA proteins may regulate MCT1. J.E.M. with guidance from H.R.C. and J.C. initiated experiments on MCT1 regulation and function. J.E.M. trained E.N.B. in the laboratory with experimental approaches on EBV-infection, drug treatments, and flow cytometry. E.N.B. and J.E.M. performed buffy coat isolations from peripheral blood, EBV infections, and early-infection treatment with AZD3965 and FACS analysis. E.N.B. performed all late-infection/LCL treatments with MCT inhibitors, phenformin and metformin, lactate measurements, Seahorse analysis, and metabolic assays. J.E.M. performed RNA-seq analysis, as well as BrdU labeling. J.M.C. performed time-course qPCR experiments as well as all P493-6 experiments. M.P. generously provided the MCT4 inhibitor VB124 and offered valuable insight on experimental design of treatments in LCLs. J.C. and H.R.C. contributed to the experimental design and execution of ^13^C6-glucose tracing. E.N.B. and M.A.L. wrote and all authors edited the manuscript.

---

**Supplementary Figure 1.**
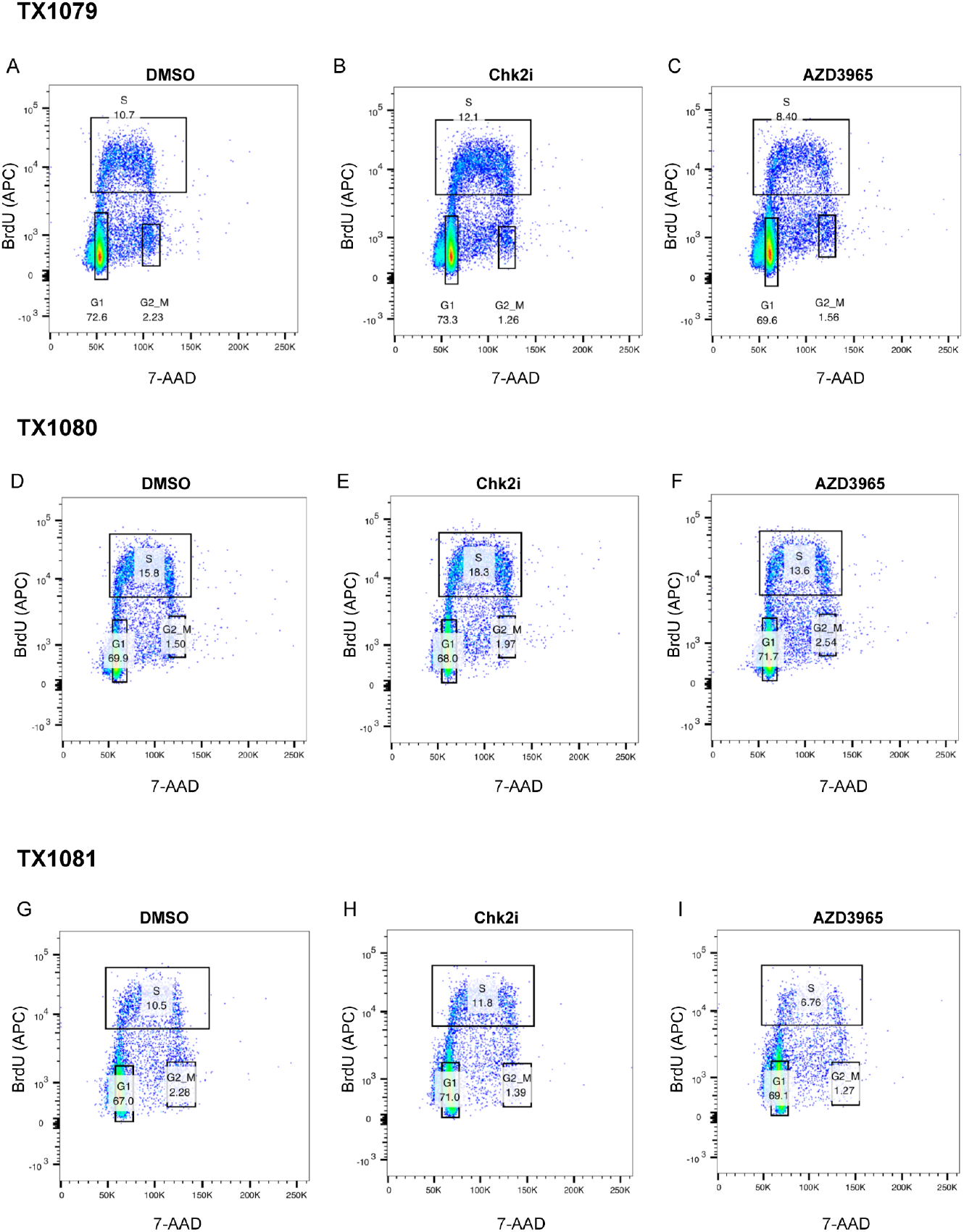
MCT1 inhibition in early EBV-infected B cells leads to a G1/S-phase growth arrest, and not cell death. ***A-I***. Three PBMC donors were treated with either DMSO, 5μM Chk2 inhibitor II, or 250nM AZD3965 for 48h. After 48h, cells were pulsed with BrdU for 2h, and cell cycle progression assessed by flow cytometry. Chk2i = positive proliferation control.

**Supplementary Figure 2.**
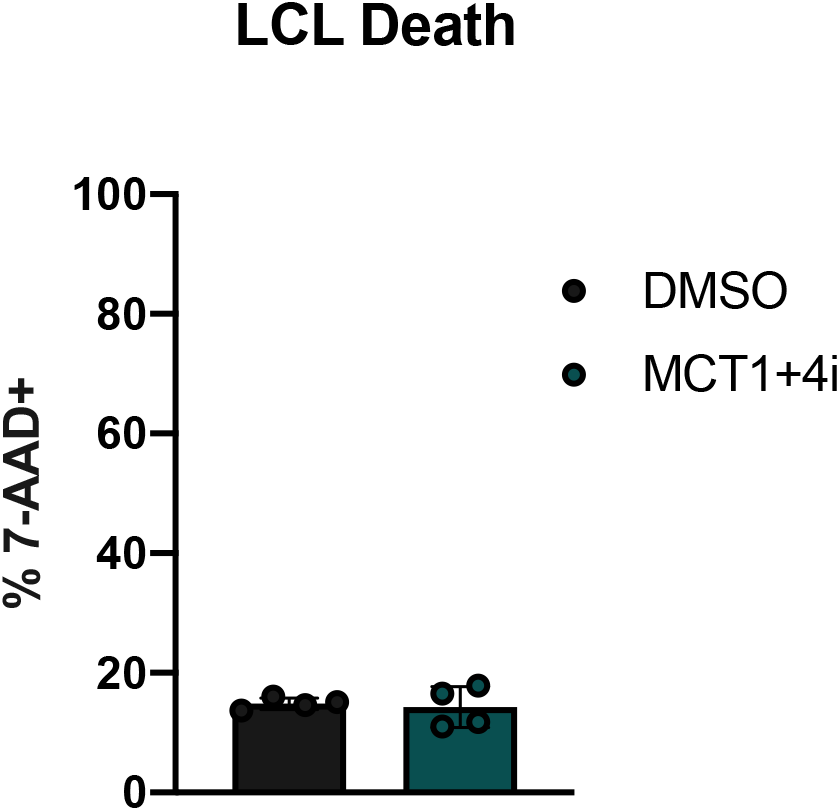
Dual MCT1/4 inhibition in LCLs does not result in cell death. Two LCL donors were treated in duplicate (n=4) with either DMSO or 1μM AZD3965 + 20μM VB124 for 72h, and 7-AAD positivity was assessed via fluorescence-activated cell sorting (FACS).

**Supplementary Figure 3.**
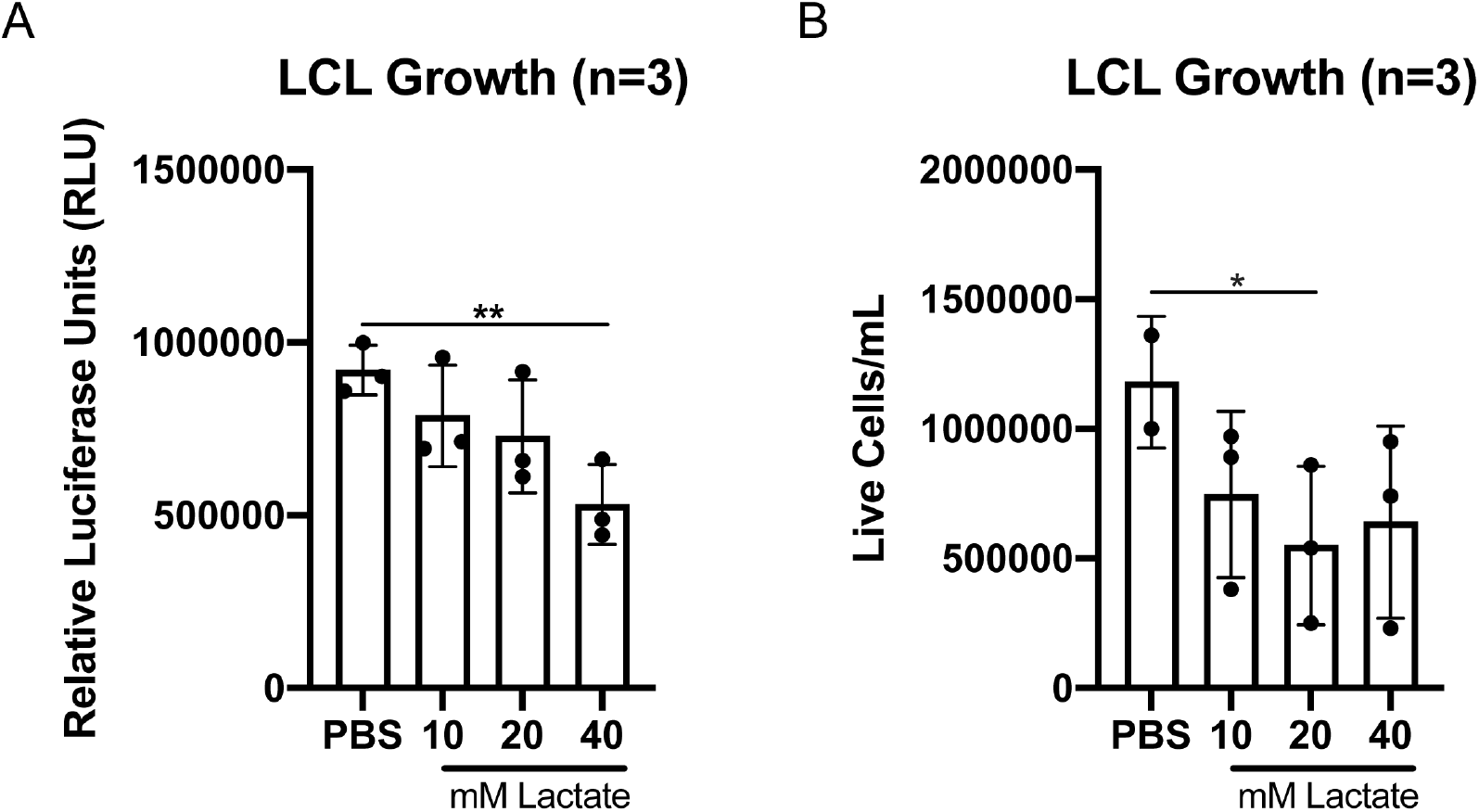
Exogenous L-lactate treatment in LCLs leads to growth arrest. LCLs were treated for 72h with varying concentrations of L-lactate. Growth was assessed by either CellTiter Glo (***A***) or trypan blue exlusion counting (***B***). Statistical significance was determined by paired t-test. *=p<0.05, **=p<0.01.

